# GlyEnzEA: reaction-informed enrichment of enzyme-centered glycan sets from glycomics data

**DOI:** 10.64898/2026.10.14.759156

**Authors:** Koichi Himori, Bingyuan Zhang, Yusuke Matsui

## Abstract

Enzyme-centered interpretation of glycomic changes requires testing whether glycans associated with particular reactions change collectively. Motif analyses and reaction databases do not directly provide enzyme-specific sets of measured glycans, and structural ambiguity or overlapping reactions can give different enzymes identical sets. A framework is needed to construct these sets and define distinguishable test hypotheses. GlyEnzEA constructs sets by matching reaction rules to candidate glycan topologies, groups enzymes sharing identical sets, and tests each set once. In an Mgat3-knockout mouse glycome, 48 enzymes yielded 16 sets; only the MGAT3 set showed significant negative enrichment after correction (Storey *q* = 0.033). Across seven HEK293 perturbations, target sets were significant in two conditions: MGAT1 and class-I mannosidase knockouts. GlyEnzEA supports enzyme-centered interpretation at the resolution of distinguishable enzymes or enzyme groups.

## 1 Introduction

Glycan profiles reflect sequential and competing enzymatic reactions, but do not uniquely identify the enzymes responsible for an observed change (Neelamegham and Liu 2011). Enzyme-centered enrichment could help interpret these profiles in terms of upstream enzymes, analogous to regulator– target analysis in transcriptomics and phosphoproteomics (Carro et al. 2010; Casado et al. 2013; Garcia-Alonso et al. 2018). Such analysis requires a glycan set for each enzyme. In glycomics, these sets must be constructed by relating reaction knowledge to the glycans measured in each dataset.

Constructing enzyme-centered sets requires reconciling structural representations and matching local reactions to observed glycans. One topology can have several identifiers, and one measured feature can have several candidate structures (Fig. 1B). Here, a feature is a source-defined quantitative measurement or measurement group, rather than necessarily a single chemical species. Reaction rules describe local transformations, whereas observed glycans may have undergone further reactions (Fig. 1E). Each enzyme-centered set therefore comprises measured features whose candidate structures are compatible with that enzyme’s rules. Different enzymes can yield identical sets, depending on the glycans observed and the structural information retained. Enrichment cannot distinguish these enzymes (Fig. 1H); collapsing identical sets defines the distinct hypotheses tested in that dataset.

**Figure 1.**
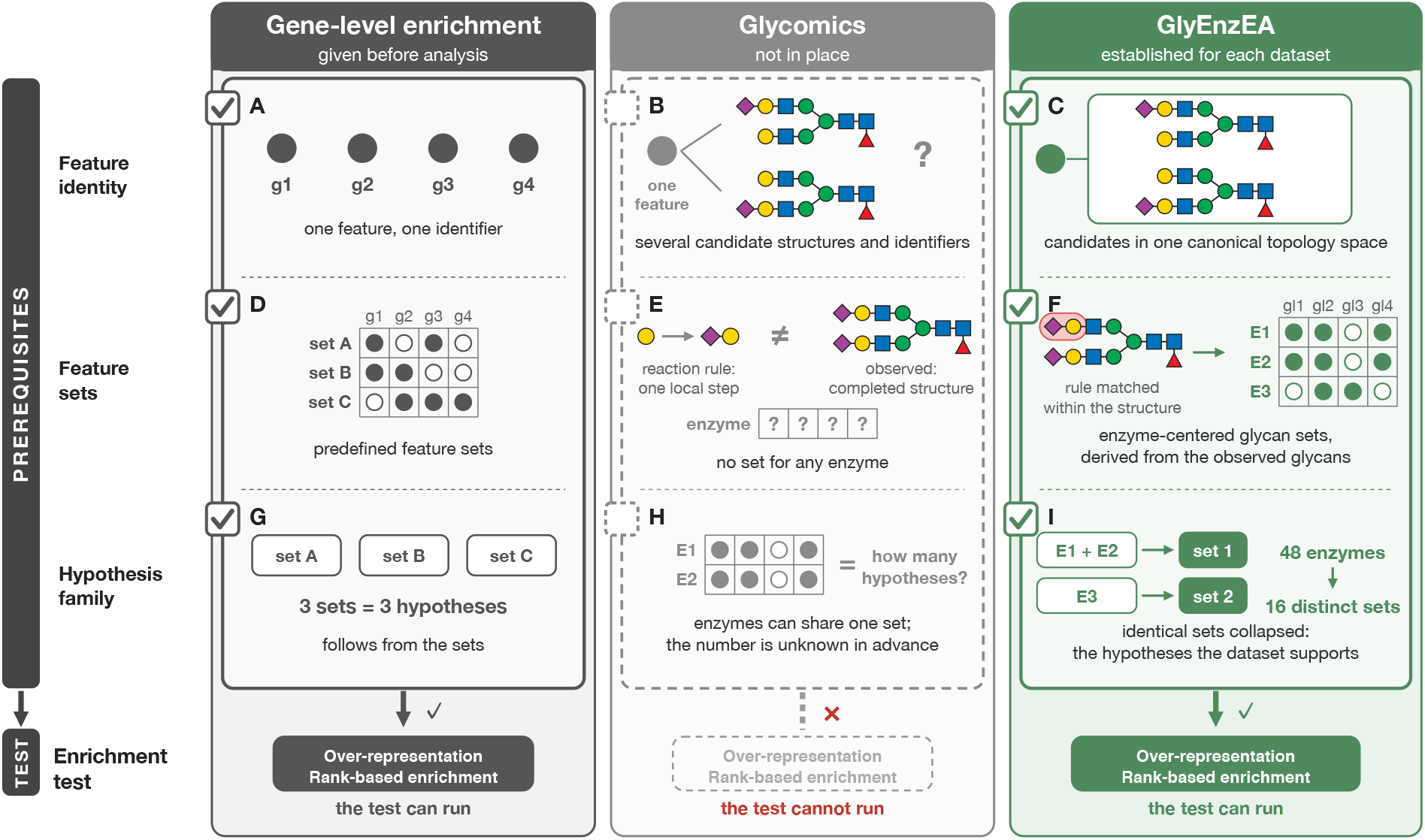
Enzyme-centered glycan sets and the hypothesis family. Columns show conventional enrichment (A,D,G), glycomics challenges (B,E,H), and GlyEnzEA (C,F,I); rows address feature identity (A–C), feature sets (D–F), and the hypothesis family (G–I). Candidate structures share a topology representation (C), structural matching defines sets (F), and identical sets are collapsed (I). Filled/open circles denote mapped/unmapped membership; question marks denote undefined membership. Matrices and +Neu5Ac are schematic; glycan examples retain source structures. In GlyEnzEA, over-representation and rank-based enrichment use one-sided Fisher’s exact tests and preranked GSEA (fgsea), respectively. **Alt text:** Diagram contrasting conventional enrichment with glycomics and showing topology matching and collapse of identical enzyme sets.

Existing resources provide enzyme–reaction knowledge (York et al. 2020; Zhou et al. 2025), while computational tools support motif-based comparison and biosynthesis-aware glycoprofile analysis (Thomes et al. 2021; Bojar and Lisacek 2022; Bao et al. 2021). The remaining task is to connect reaction knowledge to measured features to derive enzyme-centered sets and make their distinguishability explicit.

GlyEnzEA represents candidate glycan structures and reaction substrates and products in a common canonical topology space (Fig. 1C), derives set membership by substructure matching (Fig. 1F), and collapses identical sets before enrichment (Fig. 1I). The workflow tests each distinct set once using over-representation or rank-based enrichment and reports all enzymes sharing that set (Fig. 2). We evaluate the framework in an Mgat3-knockout mouse glycome and seven HEK293 perturbations. The output supports interpretation of enzyme-associated structural changes at the resolution afforded by the data, rather than directly measuring catalytic activity.

**Figure 2.**
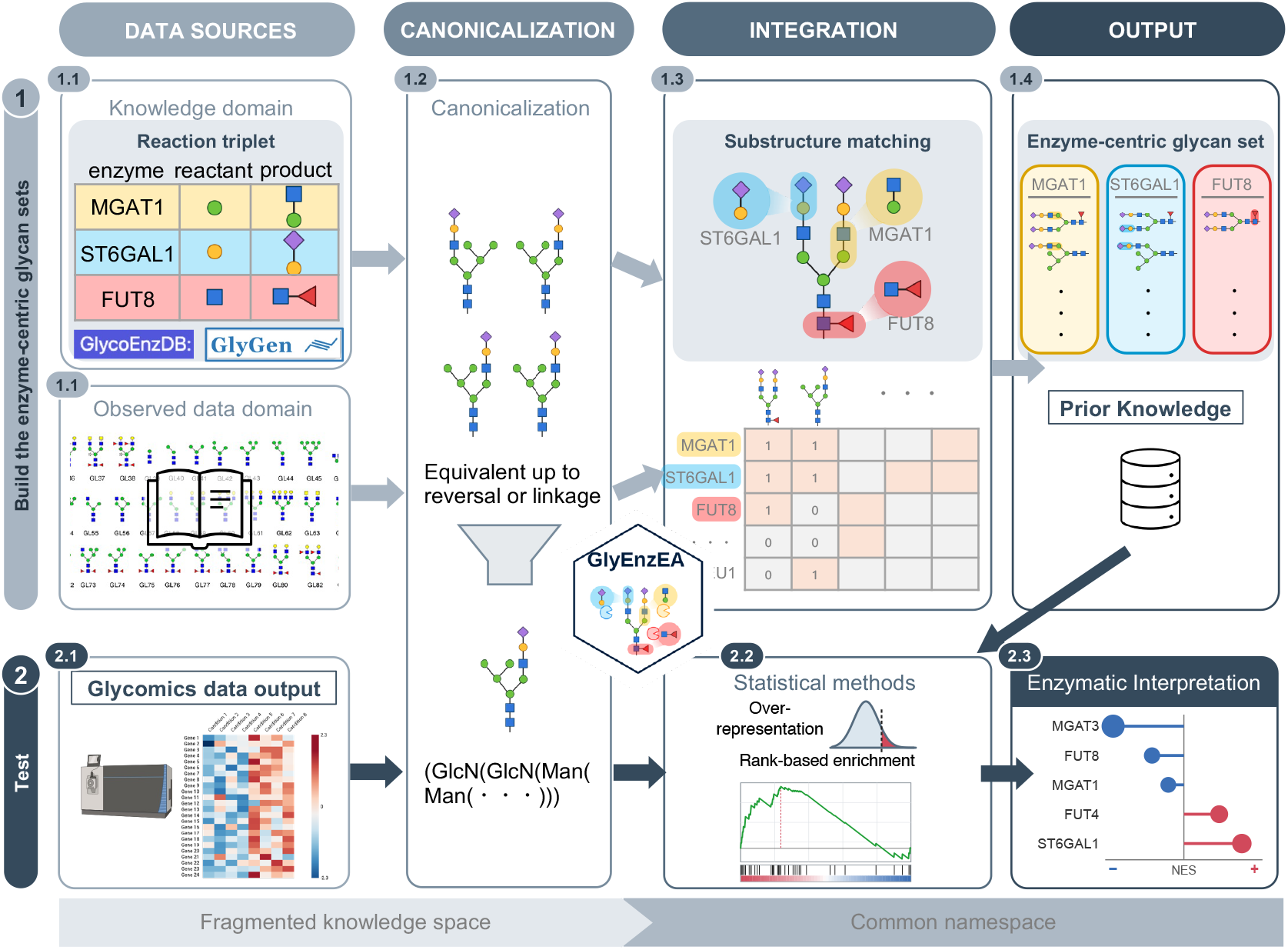
GlyEnzEA workflow in two steps. Step 1 builds enzyme-centered glycan sets: reaction triplets (enzyme, reactant, product) from GlycoEnzDB and GlyGen and observed structures (1.1) are canonicalized (1.2), reaction rules are matched by substructure containment (1.3), and the enzyme–glycan incidence matrix defines each enzyme’s set (1.4); identical sets are collapsed. Step 2 combines quantitative glycomics data (2.1), using the same canonical names, with these sets for over-representation or rank-based enrichment (2.2), using one-sided Fisher’s exact tests or preranked GSEA (fgsea), respectively, to yield enzyme-level enrichment (2.3). Colours follow MGAT1, ST6GAL1 and FUT8 from rules through matching to glycan sets; matched substructures are marked in each set member. **Alt text:** Workflow from reaction rules and glycan structures through set construction to enrichment using quantitative measurements.

## 2 Methods

### 2.1 Observations, reaction knowledge and testable hypotheses

Let 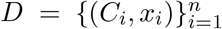 contain observations (measured features) with non-empty candidate-structure sets *C*_*i*_ and quantitative scores *x*_*i*_. Human and mouse N-glycan reactions from GlycoEnzDB and GlyGen were represented as enzyme–reactant–product triplets *r* = (*e, s, p*). We normalized carrier prefixes and notation-specific decorations, excluded unresolved sulfate/phosphate placement and deduplicated canonical topologies, obtaining 2,316 triplets for 93 enzymes.

Let *E* denote the enzymes in the reaction resource and *R*_*e*_ the rules associated with enzyme

*e*. Let *m*_*e*_(*i*) = 1 indicate that observation *i* is structurally compatible with at least one rule for *e*, as defined in Eq. (2). We define the enzyme-centered glycan set *S*_*e*_(*D*) and a dataset-specific equivalence relation between enzymes as

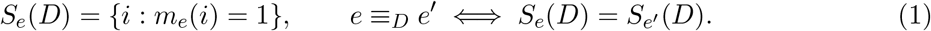

Here, *e* ≡ _*D*_ *e*^′^ means that the two enzymes map to exactly the same observations in *D*, rather than merely to the same number of observations. Such enzymes cannot be distinguished by enrichment of these sets; this does not imply equivalent biological functions.

Matches from multiple rules are combined by enzyme, then by OR across candidates within each quantitative observation. Empty glycan sets and sets spanning the mapping universe are removed at both candidate and observation stages; observation columns with no mapped enzyme are also removed from the mapping matrix. After filtering, the retained enzymes 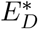 are partitioned into equivalence classes under ≡_*D*_. Each class contains enzymes sharing one identical glycan set. The collection of these classes, 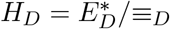, defines the hypothesis family: each distinct set is tested once, and all enzymes in its class are reported.

For rank-based enrichment, each measured feature with valid structural annotation and a finite score is ranked individually. Enzymes sharing an identical glycan set are tested together, whereas distinct measured features retain their individual scores and are not merged even when they map to the same enzymes. Filtering order and missing-value handling are detailed in Supplementary Methods. Set sizes, Jaccard similarities and binary *φ* coefficients describe remaining overlap.

### 2.2 Topology canonicalization and structural matching

To compare enzyme reaction rules and observed glycans across notations, we represent reactants, products and candidate glycans as rooted, monosaccharide-labeled trees. The map *ϕ* retains monosaccharide labels and branching topology but omits anomer and linkage position; glycans differing in these details may share one topology. AHU-style recursive subtree encoding removes branch-order differences by canonicalizing child order (Aho et al. 1974). The topoSeq string encodes the canonical tree using monosaccharide names and parentheses, assigning identical strings to identical labeled topologies (Supplementary Methods S3.1). Duplicate candidate topologies are merged within each observation, but measured-feature abundances are not allocated among candidates.

Next, we match candidate glycans to reaction rules to assign measured features to enzyme-centered glycan sets. Observed glycans may carry modifications from other reactions beyond the local structures in a rule, so we test substructure containment rather than whole-structure identity (Fig. 1E,F).

Let *g* ∈*C*_*i*_ denote a candidate structure for observation *i*. The relation *a* ⪯ *g* means that *a* is contained in *g*, preserving monosaccharide labels and parent-to-child edge directions. Transferase rules require containment of both the canonical reactant and product, *ϕ*(*s*) ⪯ *ϕ*(*g*) and *ϕ*(*p*) ⪯ *ϕ*(*g*). For GlycoEnzDB Glycosidase rules, a candidate glycan must contain the product substructure but not the reactant substructure, selecting structures compatible with sugar-residue cleavage. Source- and class-specific criteria are given in Supplementary Methods S3.

A measured feature belongs to enzyme *e*’s glycan set if any candidate structure satisfies the matching criteria of any rule for *e*. The membership indicator is

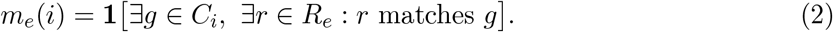

Containment is tested on the trees represented by topoSeq, rather than on the strings, using VF2 implemented in igraph (Cordella et al. 2004; Csardi and Nepusz 2006). Reactant and product structures are matched independently without requiring a shared reaction-center alignment, so they may match different branches of a candidate glycan. Root-to-root alignment and cellular compartment are also unconstrained. Thus, set membership indicates structural compatibility, not evidence of an actual reaction or biosynthetic path.

### 2.3 Ranking, enrichment and robustness assessment

Features are ranked by log-ratios of relative abundances (percentages) in perturbation 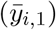 and control 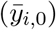 conditions:

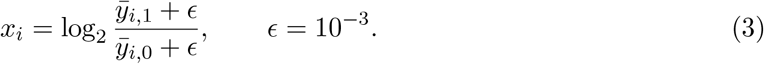

The 0.001-percentage-point pseudocount handles zeros. HEK293 uses source-reported condition means; mouse values are aggregated by source annotation. Each feature contributes one score regardless of candidate count. Weighted enrichment depends on ranks and magnitudes, so the pseudocount can affect results without changing ranks.

Over-representation analysis tests whether glycan-set members are over-represented among increased or lost features. We apply one-sided Fisher’s exact tests (ORA) separately to source-annotated increased and lost lists, using the union of valid structurally annotated feature identifiers and glycan-set identifiers as background.

Rank-based enrichment tests whether set members concentrate at the high or low end of the scores in Eq. (3). We apply preranked gene set enrichment analysis (GSEA) using fgsea (Korotkevich et al. 2019), with a weighted Kolmogorov–Smirnov-type score and adaptive multilevel *P* -value estimation. The minimum set size is one; detailed settings are in Supplementary Methods S4. These *P* -values assess enrichment within the ranked features, not biological-replicate variation or catalytic activity. Single-feature sets are recorded but do not support confirmatory claims.

Correction covers each hypothesis family: 16 distinct mouse glycan sets or 22 per HEK293 perturbation. Storey *q*-values use qvalue with bootstrap estimation of the true-null proportion *π*_0_; estimation failure triggers Benjamini–Hochberg (BH) adjustment. We denote the adopted corrected value by *q*, with significance at *q <* 0.05. Tables S1 and S2 report the correction used and BH-adjusted values. This correction does not control false discoveries across perturbations.

Mouse MGAT3 enrichment is compared with size-matched random sets, enzyme-label reassignment and degree-preserving curveball matrices (Strona et al. 2014), assessing set size, position among existing sets and enrichment conditional on enzyme and feature degrees, respectively. The 1,000 curveball matrices give empirical *P* -value resolution of approximately 0.001. Label reassignment evaluates MGAT3’s enrichment-score rank among the 16 sets. Sensitivity analyses vary nPermSimple from 10^3^ to 10^5^ over five random seeds (Supplementary Methods S4).

### 2.4 Validation design and evaluation criteria

We evaluate detection of glycan-set changes associated with known gene deletions using Mgat3-knockout mouse and HEK293 perturbation data. Target sets and expected directions are defined from the affected reactions and structural mapping. Detection in the expected direction requires both primary *q <* 0.05 and the expected sign of the normalized enrichment score (NES). Unadjusted *P* -value ranks supplement the assessment of non-significant target sets. Enzymes sharing a set cannot be distinguished individually.

In the mouse brain dataset (Nakano et al. 2019), we rank 33 of 34 measured features after excluding feature 10d, which lacks structural annotation. Multiple candidates are retained within each feature, while measurements reported separately in the source remain distinct (Supplementary Methods S1). Mgat3 knockout is the positive control, with expected negative enrichment of the MGAT3 set. The FUT8 set is a negative comparison based on unchanged core-fucose-specific PhoSL binding; non-significance alone does not establish equivalence. Terminal sialylation and antennary fucosylation provide secondary, direction-only assessments of the corresponding sets.

Applicability across perturbations is assessed in seven HEK293 gene-deletion conditions (Huang et al. 2021). Excluding one unannotated feature leaves 40 of 41 measured features. Candidates are expanded from multiple structure identifiers and ambiguous linkage information, and all conditions share one enzyme–glycan mapping (Fig. 5; Supplementary Methods S5). To interpret non-detection, we also consider WT target-glycan abundance, using values below 2% as a low-abundance guide (MGAT5: 0.6%). This is neither an exclusion criterion nor an empirically established detection limit. SLC35C1 deletion, which lacks an enzyme reaction rule, is assessed indirectly through the FUT8 set as a readout of fucosylation.

To evaluate mapping coverage and processing consistency, we use the glycan structure library for the glycoproteomics software Glyco-Decipher (Fang et al. 2022) as a reference collection, canonicalized independently of the mouse data. Coverage uses all original records as the denominator, including conversion failures. Consistency is the fraction of common enzyme–topology pairs with identical assignments in the library and mouse matrices, excluding pairs with missing values. Both mapped–mapped and unmapped–unmapped pairs count as agreement. This measures consistency under shared reaction rules, not independent biological mapping accuracy.

## 3 Results

### 3.1 Mapping coverage and testable glycan sets

To assess enzyme mapping across diverse glycan structures, we used an existing glycan structure library as a reference (Fang et al. 2022). Of 10,936 records, 8,098 were converted into 3,299 distinct topologies, all of which mapped to at least one enzyme. The mappings involved 84 enzymes in total, with 2–50 enzymes per topology.

We next examined which enzymes could be distinguished for testing in measured data. The 33 measured features in the mouse dataset were represented by 97 candidate topologies and mapped to 48 enzymes. Collapsing enzymes with identical glycan sets yielded 16 distinct sets (Fig. 3). Six, including MGAT3 and FUT8, each corresponded to a single enzyme; the remaining ten were shared by 2–10 enzymes. Each set contained 1–29 measured features, and each of the 16 sets was tested once.

**Figure 3.**
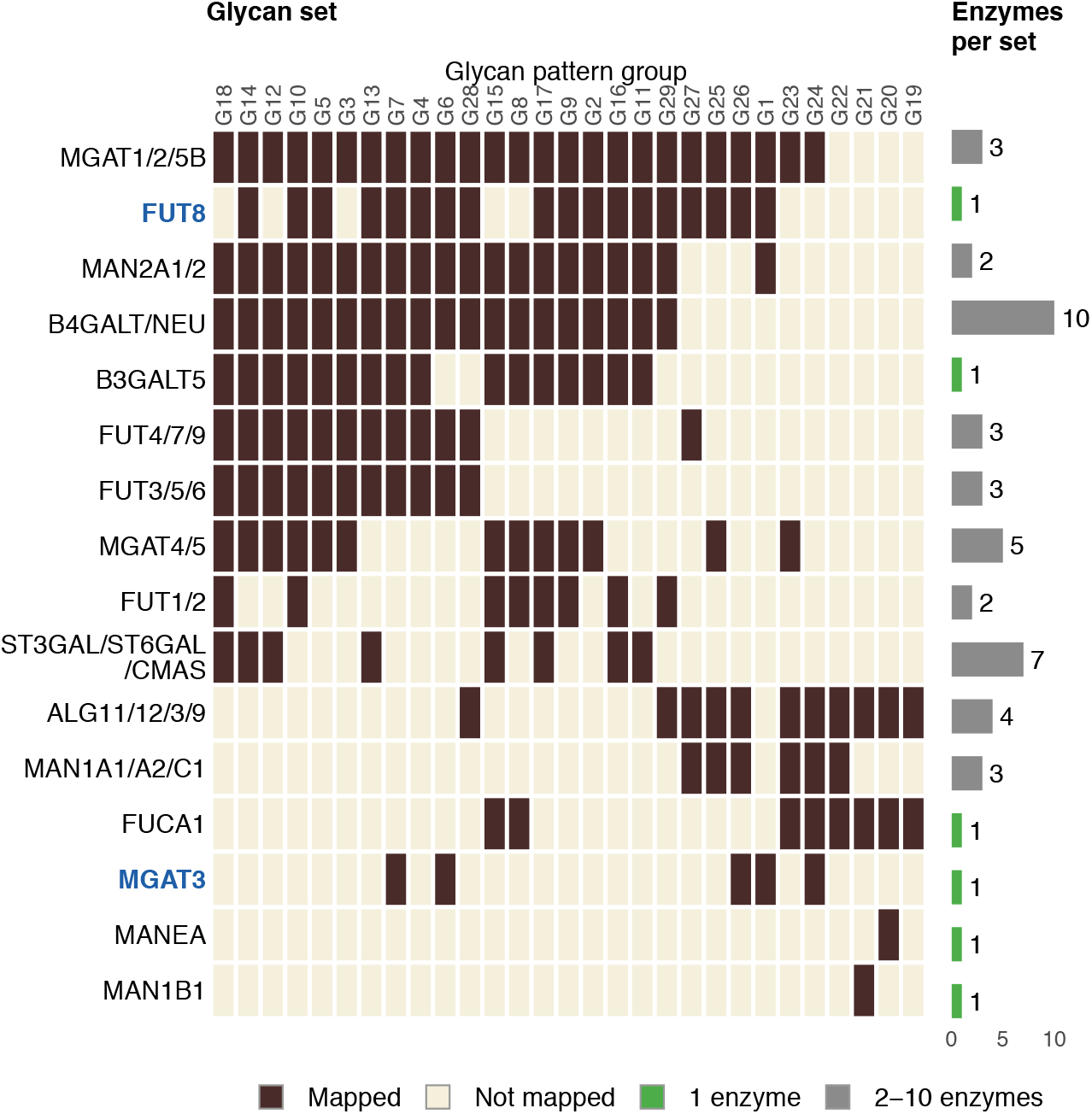
Dataset-specific glycan sets: 48 enzymes yield 16 distinct sets. Rows show 16 glycan set membership patterns in one continuous matrix; columns group 97 candidate topologies into 29 identical enzyme-mapping patterns for display, distinct from the 33 measured features used for enrichment. Dark cells indicate mappings. Bars count enzymes sharing each row, not mapped glycan groups: six single-enzyme sets (green) and ten shared sets (grey). The MGAT3 and FUT8 glycan sets are highlighted. Glycan examples are in Supplementary Fig. S1. **Alt text:** Membership matrix shows sixteen distinct sets for forty-eight enzymes, including six single-enzyme and ten shared sets.

Shared sets indicate that enzymes cannot be distinguished with these data and this structural representation. For example, sialyltransferase-related enzymes shared one set, while B4GALT1–5, NEU1–4 and B3GALT2 shared another. Shared substrates, missing distinguishing glycans and omitted linkage information can produce identical sets without implying functional interchangeability. Overlap also remained between distinct sets after identical sets were collapsed: sets associated with later biosynthetic steps had Jaccard indices of 0.79–0.96.

Comparison with the library identified 66 shared topologies among the 97 mouse candidates (68%). Mapping presence or absence agreed for all 3,168 common enzyme–topology pairs, confirming processing consistency under shared reaction rules.

### 3.2 Detection of glycan-set changes following MGAT3 loss

In Mgat3-knockout mice, only the MGAT3 glycan set was significantly enriched after multipletesting correction (NES = −1.60, unadjusted *P* = 0.003, Storey *q* = 0.033; Fig. 4C). All seven measured features in this set included a candidate structure bearing bisecting GlcNAc and disappeared in knockout mice (Fig. 4A). They therefore clustered at the decreasing end of the ranking (Fig. 4B). Over-representation analysis of lost features was also significant (*P* = 7.7 × 10^−5^), and the rank-based result remained significant under BH adjustment (adjusted *P* = 0.049).

**Figure 4.**
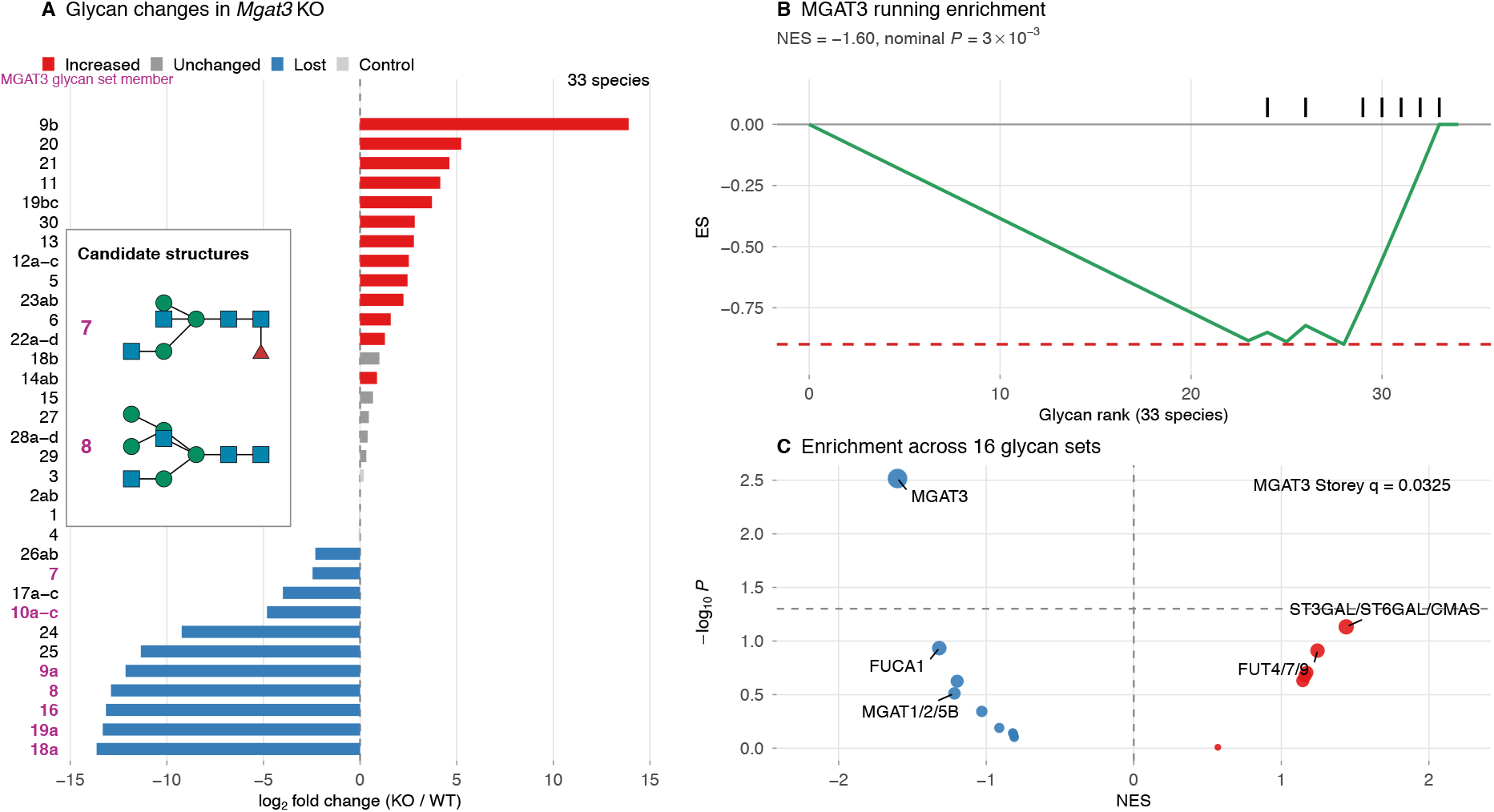
Glycan set changes following MGAT3 loss. (A) log_2_ fold changes for the 33 measured features used for ranking, with original increased/unchanged/lost/control colours. Magenta marks identify seven features in the MGAT3 glycan set, all lost; two candidate structures appear in the inset. (B) Running enrichment across 33 ranked measured features; ticks mark MGAT3 glycan set members and the dashed line marks minimum ES (−0.90). (C) NES versus nominal *P* for 16 glycan sets; blue/red denote negative/positive NES, size indicates −log_10_ *P* and the dashed line denotes nominal *P* = 0.05. Only the MGAT3 glycan set is significant after correction (NES = −1.60, Storey *q* = 0.033). See Supplementary Fig. S2 for degree-preserving null validation. **Alt text:** Fold changes, running enrichment and set scores show depletion of the MGAT3 set after Mgat3 knockout.

Randomization preserving mapping counts per enzyme and per feature supported this result (empirical *P* = 0.001; Supplementary Fig. S2). This value corresponds to the resolution limit of 1,000 randomizations. Enzyme-label reassignment ranked MGAT3 first among the 16 sets, although this assessment has a resolution of only 1/16. Significance under Storey’s method was retained when minimum set size varied from 1 to 3 and nPermSimple from 10^3^ to 10^5^. The significance decision under BH adjustment depended on the computational settings.

Sets associated with terminal sialylation and antennary fucosylation shifted in the direction of the increased terminal modifications reported in knockout mice (Nakano et al. 2019). The ST3GAL/ST6GAL/CMAS, FUT4/7/9 and FUT3/5/6 sets had NES values of 1.44, 1.25 and 1.16, respectively, but none was significant (unadjusted *P* = 0.074, 0.12 and 0.21). The FUT8 set also showed no significant enrichment (NES = −0.81, unadjusted *P* = 0.77). This is compatible with the reported unchanged core-fucose-specific PhoSL binding, but does not establish unchanged core fucosylation. Thus, the main statistically supported result was depletion of the MGAT3 set; terminal modifications showed directional concordance.

### 3.3 Detection and interpretation of HEK293 target sets

We assessed applicability across biosynthetic steps using seven HEK293 gene-deletion conditions (Huang et al. 2021). The 40 analyzed measured features mapped to 60 enzymes and 22 distinct glycan sets. Testing these 22 sets in each condition identified significant target-set enrichment in two conditions, MGAT1 and class-I mannosidase knockout; both were also significant under BH adjustment (Fig. 5; Supplementary Table S2). Target ranks by unadjusted *P* value ranged from 2 to 16 (median 6), providing supplementary context for non-significant targets.

**Figure 5.**
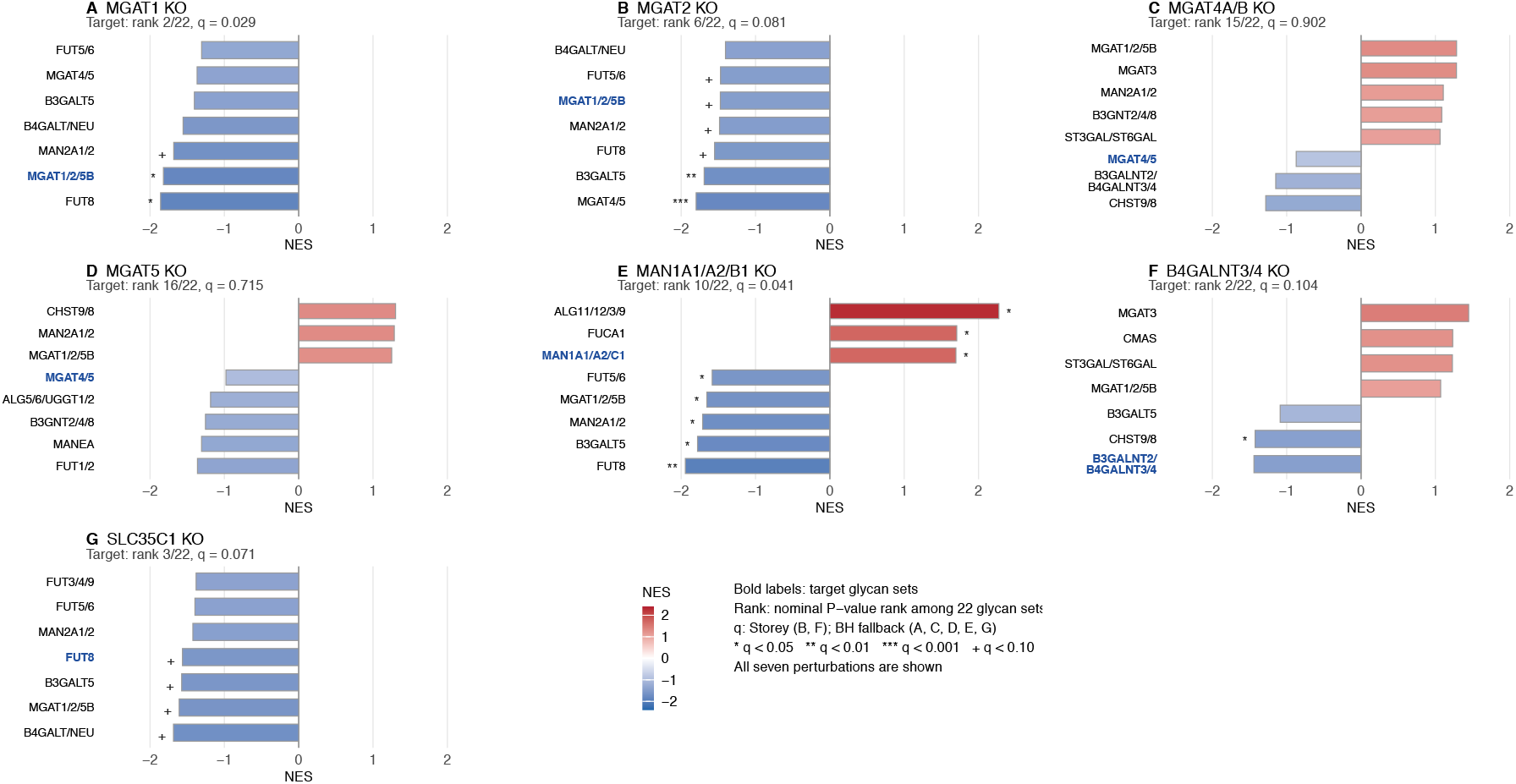
Glycan set results across seven HEK293 perturbations. (A–G) Selected glycan set NES for the indicated knockouts; bold labels identify target sets. Headings give target nominal-*P* rank among 22 glycan sets and within-perturbation *q* (Storey in B and F; BH fallback in A, C–E and G). Shared scales and significance symbols are defined in the legend; complete results are in Supplementary Table S2. ST3GAL/ST6GAL and B3GALNT2/B4GALNT3/4 abbreviate labels for shared glycan sets. In (E), the target glycan set is associated with MAN1C1 whereas the deletion includes MAN1B1. **Alt text:** Seven perturbation panels show significant target-set enrichment only for MGAT1 and class-I mannosidase knockouts.

In MGAT1 knockout, the MGAT1/2/5B set showed significant negative enrichment (NES = −1.82, unadjusted *P* = 0.0026, *q* = 0.029). Of its 29 measured features, 27 decreased, and combined relative abundance fell from 15.3% to 0.56%. In class-I mannosidase knockout, the target set showed significant positive enrichment (NES = 1.70, *q* = 0.041). Man6–Man8 intermediates accumulated while Man5, produced after mannose removal, decreased. The positive NES was therefore consistent with intermediate accumulation following blocked sequential mannose removal.

In the other conditions, target sets showed directional or ranking patterns without reaching significance. In MGAT2 knockout, the same shared set as in MGAT1 knockout had a negative NES (NES = −1.47, *q* = 0.081; unadjusted-*P* rank 6). In B4GALNT3/4 knockout, the target had the most negative NES among all sets but was non-significant (NES = −1.44, *q* = 0.104; rank 2) ; only one of two measured features bearing LacdiNAc disappeared. In SLC35C1 knockout, the FUT8 set, used to assess effects on fucosylation indirectly, was also non-significant (NES = −1.56, *q* = 0.071; rank 3). All five fucosyltransferase-associated sets nevertheless had negative NES values.

The shared MGAT4/5 set was non-significant in MGAT4A/B and MGAT5 knockouts (unadjusted-*P* ranks 15 and 16, respectively). Omitting *β*1, 4 and *β*1, 6 linkage information merges the tri-antennary products of these enzyme groups into the same topology. This shared set cannot distinguish glycan changes specific to either group.

## 4 Discussion

GlyEnzEA constructs enzyme-centered glycan sets from reaction knowledge and tests enrichment at the resolution supported by the observations and structural representation. The reduction from 48 enzymes to 16 distinct sets in the mouse dataset shows that enzyme counts differ from the number of distinguishable test hypotheses (Fig. 3). Grouping enzymes with identical sets avoids duplicate tests of the same observations and makes explicit whether a result corresponds to one enzyme or is shared by several. This distinction defines the scope of enzyme-related hypotheses drawn from enrichment results.

In evaluations using known gene deletions, the mouse MGAT3 set and the target sets in two of seven HEK293 perturbations showed significant enrichment (Figs. 4 and 5). These findings demonstrate detection of glycan sets associated with known perturbations; performance in identifying unknown causal enzymes requires separate evaluation. Mouse sets associated with terminal sialylation and antennary fucosylation shifted in the direction of previously reported increases in terminal modifications, but were not significant (Nakano et al. 2019). Non-significance of the FUT8 set likewise does not establish unchanged core fucosylation. These findings provide hypotheses for investigating secondary changes.

Enrichment direction and non-detection require interpretation against member glycans and the affected reaction. Positive enrichment after class-I mannosidase loss was consistent with Man6– Man8 intermediate accumulation and does not indicate increased enzyme activity. The shared MGAT4/5 set illustrates a structural limitation: omitting linkage positions merges products of both enzyme groups into the same topology. Low target-glycan abundance or insufficient observations may also contribute to non-detection. Non-significance alone therefore establishes neither absence of a reaction-level effect nor the cause of non-detection.

Among existing glycan analysis tools, Glycowork supports motif analysis and GlyCompare organizes glycan profiles through biosynthetic relationships (Thomes et al. 2021; Bao et al. 2021). GlyEnzEA maps observed glycans to enzyme reactions and tests sets defined by those mappings. These methods support different interpretations, so comparing detection counts alone is insufficient. Comparisons with motif-based analyses and linkage-preserving matching require common evaluation tasks and datasets.

Structural mapping is limited by reaction-resource coverage and matching criteria. Omitting anomers and linkage positions removes distinctions needed to discriminate enzymes. Independent reactant and product matching can satisfy the criteria on different branches, and tissue or compartment constraints are not considered. Including a feature whenever any candidate matches can also broaden membership when annotations are ambiguous. Set membership therefore represents structural compatibility with a reaction. Agreement with the reference library confirms processing consistency under shared rules, rather than independently validating biological accuracy.

Statistical interpretation also has limits. Glycans remain shared between distinct sets after identical sets are collapsed, leaving dependence among test results. Estimating the null proportion for Storey’s method may be unstable in hypothesis families of only 16–22 sets; comparisons with BH adjustment and alternative computational settings assess this sensitivity. Relative-abundance scores depend on normalization and pseudocounts, and rank-based enrichment *P* values do not assess variation among biological replicates. Reproducibility of detected sets therefore requires validation in independent data.

Future work should improve set specificity through linkage-preserving matching and reaction-center alignment, and examine how substrates and products are represented in sequential reactions. Beyond extending validation with known deletions, evaluation should establish how well the method narrows candidate enzyme groups and reaction steps when the causal perturbation is unknown. This would support using glycan changes, including those associated with disease, to generate hypotheses about enzyme reactions for experimental validation. The reaction-based mapping and set-construction strategy could also be extended to site-specific N-glycoproteomics when candidate glycan structures are available. This application was not evaluated here and would require appropriate normalization, treatment of dependence among glycopeptide measurements, and dedicated validation.

## Data and code availability

R code and workflow: https://github.com/matsui-lab/GlyEnzEA. Code, test data and frozen inputs are archived at https://doi.org/10.5281/zenodo.23345861 (Supplementary Methods S6). Original code is MIT-licensed; upstream data retain their source rights and conditions.

## Supplementary information

Supplementary information accompanies this manuscript and includes Supplementary Methods S1–S7, Tables S1–S2 and Figures S1–S2.

## Author contributions

Koichi Himori: methodology, software, formal analysis, validation and writing—original draft.

Bingyuan Zhang: methodology and formal analysis.

Yusuke Matsui: conceptualization, methodology, validation, writing—original draft and supervision.

## Acknowledgements

The authors thank the developers and curators of GlycoEnzDB, GlyGen, Glyco-Decipher and the open-source R packages used in this study.

## Use of generative AI

Generative AI was used for English-language editing, translation and assistance with code writing. Codex (OpenAI) also assisted manuscript formatting and submission preparation. AI assistance is disclosed here as part of the research and writing workflow; responsibility for the manuscript and code rests with the authors.

## References

Aho A V, Hopcroft JE, Ullman JD. The Design and Analysis of Computer Algorithms. Reading (MA): Addison-Wesley, 1974.

Bao B, Kellman BP, Chiang A WT et al. Correcting for sparsity and interdependence in glycomics by accounting for glycan biosynthesis. Nat Commun 2021;12:4988.

Benjamini Y, Hochberg Y. Controlling the false discovery rate: a practical and powerful approach to multiple testing. J R Stat Soc Ser B 1995;57:289–300.

Bojar D, Lisacek F. Glycoinformatics in the artificial intelligence era. Chem Rev 2022;122:15971–88.

Carro MS, Lim WK, Alvarez MJ et al. The transcriptional network for mesenchymal transformation of brain tumours. Nature 2010;463:318–25.

Casado P, Rodriguez-Prados JC, Cosulich SC et al. Kinase-substrate enrichment analysis provides insights into the heterogeneity of signaling pathway activation in leukemia cells. Sci Signal 2013;6:rs6.

Cordella LP, Foggia P, Sansone C et al. A (sub)graph isomorphism algorithm for matching large graphs. IEEE Trans Pattern Anal Mach Intell 2004;26:1367–72.

Csardi G, Nepusz T. The igraph software package for complex network research. InterJournal Complex Systems 2006;1695:1–9.

Fang Z, Qin H, Mao J et al. Glyco-Decipher enables glycan database-independent peptide matching and in-depth characterization of site-specific N-glycosylation. Nat Commun 2022;13:1900.

Garcia-Alonso L, Iorio F, Matchan A et al. Transcription factor activities enhance markers of drug sensitivity in cancer. Cancer Res 2018;78:769–80.

Huang Y, Aoki K, Akase S et al. Global mapping of glycosylation pathways in human-derived cells. Dev Cell 2021;56:1195–209.e7.

Korotkevich G, Sukhov V, Sergushichev A. Fast gene set enrichment analysis. bioRxiv 2019. 10.1101/060012, preprint: not peer reviewed.

Nakano M, Mishra SK, Tokoro Y et al. Bisecting GlcNAc is a general suppressor of terminal modification of N-glycan. Mol Cell Proteomics 2019;18:2044–57.

Neelamegham S, Liu G. Systems glycobiology: biochemical reaction networks regulating glycan structure and function. Glycobiology 2011;21:1541–53.

Storey JD. A direct approach to false discovery rates. J R Stat Soc Ser B 2002;64:479–98.

Strona G, Nappo D, Boccacci F et al. A fast and unbiased procedure to randomize ecological binary matrices with fixed row and column totals. Nat Commun 2014;5:4114.

Thomes L, Burkholz R, Bojar D. Glycowork: a Python package for glycan data science and machine learning. Glycobiology 2021;31:1240–4.

York WS, Mazumder R, Ranzinger R et al. GlyGen: computational and informatics resources for glycoscience. Glycobiology 2020;30:72–3.

Zhou Y, Ghosh V, Venkatesan S et al. GlycoEnzDB: a database of enzymes involved in human glycosylation. Glycobiology 2025;35:cwaf074.

