## Supplementary Information for "GlyEnzEA: reaction-informed enrichment of enzyme-centered glycan sets from glycomics data"


| | |
|---|---|
| Supplementary Methods S1–S6 | pages 1–2 |
| Table S1. Glycan sets of the Mgat3-knockout mouse glycome: composition, GSEA, correction, null checks | page 3 |
| Table S2. GSEA of all 22 glycan sets in the seven HEK293 perturbations | page 4 |
| Figure S1. Candidate structures in each display column of Figure 3 | page 7 |
| Figure S2. Null checks for the mouse enrichment results | page 9 |
| S7. Use of generative AI and author contributions | page 10 |


### Supplementary Methods

#### S1. Mouse glycome input and measured features

The Mgat3-knockout mouse brain N-glycome (Nakano et al., 2019) was used as received from the Kizuka laboratory: one row per measured feature, numbered 1–30 with letter suffixes for isomers of one composition, with relative abundances (%) in Mgat3$^{+/+}$ and Mgat3$^{-/-}$ brain and, for most features, one or more candidate structures (GlyTouCan identifiers). Suffixed rows of one composition were pooled into one measured feature unless the paper itself reports them as separate measurements (9a/9b, 18a/18b, 19a/19bc). Species 10d, which has no structure in the paper and increases in the knockout while 10a–10c disappear, is kept apart from 10a–c; having no structure, it is not mapped and not ranked. This gives 34 measured features with 98 candidate structures, of which 33 features enter the ranking. The relative abundances are normalized to the total of oligomannose-type N-glycans and are therefore not a closed composition; scores are $x_i = \log_2[(\bar{y}_{i,1}+\epsilon)/(\bar{y}_{i,0}+\epsilon)]$ with $\epsilon = 10^{-3}$ percentage points, one score per measured feature regardless of the number of candidates.

#### S2. Filtering order and missing values

Structural matching (Methods 2.2) first produces a binary matrix of enzymes by candidate topologies (93 enzymes × 97 topologies in the mouse dataset). Filtering is applied in the following order. (i) At the candidate level, enzymes that match every candidate topology are removed, then enzymes that match none. (ii) Candidate columns are combined into measured features by OR, i.e. a feature is in an enzyme's glycan set if any of its candidates matches. (iii) At the feature level, feature columns to which no enzyme maps are removed, then enzymes that map to every remaining feature, then enzymes that map to none. Feature-level filtering is repeated after OR aggregation because a set may span all retained features even when it did not span all candidate topologies. In the mouse dataset this leaves 48 enzymes and 33 features; identical rows are then collapsed into the 16 distinct glycan sets of the hypothesis family. Missing values arise only from features without a structure (10d), which are dropped before matching; scores are never missing because of the pseudocount. The ranked universe is the set of features with at least one structure and a finite score (33), and features are not merged when their enzyme-mapping patterns coincide.

#### S3. Reaction rules and matching criteria

Human and mouse N-glycan reactions were taken from GlycoEnzDB and GlyGen and represented as enzyme–reactant–product triplets. Carrier prefixes and notation-specific decorations were normalized, reactions with unresolved sulfate or phosphate placement were excluded, and canonical topologies were deduplicated, giving 2,316 triplets for 93 enzymes. Whether a rule $r = (e, s, p)$ matches an observed structure $g$ is decided by subgraph containment ($\preceq$, VF2 in igraph) of the canonical topologies, with the criterion depending on the source and class of the rule:

| Source | Enzyme class | Criterion |
|---|---|---|
| GlycoEnzDB | Glycosyltransferase | $\phi(s) \preceq \phi(g)$ and $\phi(p) \preceq \phi(g)$ |
| GlycoEnzDB | Glycosidase | $\phi(p) \preceq \phi(g)$ and not $\phi(s) \preceq \phi(g)$ |
| GlycoEnzDB | (empty) or N_linked | as Glycosyltransferase (joint reactant–product criterion) |
| GlyGen | all | as Glycosyltransferase (joint reactant–product criterion) |

Reactant and product containment are evaluated independently, without a shared reaction-centre alignment; neither root-to-root alignment nor compartment is imposed. Matches from several rules of one enzyme are combined by OR.

#### S3.1. Topology encoding and substructure comparison

Each canonical topology is a rooted tree with monosaccharide-labeled nodes and edges directed from parent to child. Anomer and linkage position are omitted, so equality refers to the retained topology, not necessarily to the same chemical structure.

To determine a reproducible branch order, AHU-style labels are computed recursively from leaves to root (Aho et al., 1974). A leaf receives its monosaccharide label followed by `()`; an internal node receives its label followed by parentheses enclosing its children's complete AHU labels sorted lexicographically. For serialization, child subtrees are visited in descending node count, with ties resolved by lexicographic order of their complete AHU labels, including parentheses.

The resulting topoSeq notation encodes each node as `(label child1 child2 ...)`, omitting spaces; a leaf is `(label)`. Parentheses delimit subtrees. For example, with abstract labels A, B and C, `(A(B)(C))` represents root A with children B and C. Reversing the input branches to `(A(C)(B))` gives the same canonical string, whereas `(A(B(C)))` represents a distinct chain. The internal AHU labels determine ordering; topoSeq is the final tree serialization.

For matching, topoSeq strings are parsed into trees. Monosaccharide labels receive shared integer vertex colors, and VF2 (Cordella et al., 2004) in igraph (Csardi and Nepusz, 2006) tests an injective embedding that preserves node labels and directed parent–child edges. Matching is performed on graphs, not by substring search. The root of the reaction pattern need not map to the glycan root; additional branches outside the matched pattern are allowed.

#### S4. Enrichment, correction and null checks

ORA uses one-sided Fisher tests on the source-annotated increased and lost lists against the union of valid annotated features and set members. Preranked GSEA uses fgsea (adaptive multilevel estimation, nPermSimple = 50,000, minSize = 1, maxSize = 500 for the mouse dataset and no upper limit for HEK293, eps = 0 and seed 42 for HEK293). The primary correction is the Storey $q$-value (qvalue, bootstrap estimation of $\pi_0$) within each hypothesis family (16 sets in the mouse dataset, 22 per HEK293 perturbation); when $\pi_0$ cannot be estimated the Benjamini–Hochberg adjustment is used instead, and Tables S1 and S2 state which was used. For the mouse dataset Storey estimation succeeded; for HEK293 it succeeded in two of the seven perturbations (MGAT2 and B4GALNT3/4). Three null checks accompany the mouse result: 1,000 degree-preserving curveball randomizations of the enzyme–feature matrix (empirical resolution 0.001), reassignment of the 16 set labels (resolution 1/16), and size-matched random sets. Sensitivity to minSize (1–3) and to nPermSimple ($10^3$–$10^5$, five seeds each) is reported in Table S1 and Figure S2.

#### S5. HEK293 perturbation panel

From the N-glycan section of Table S4 of Huang et al. (2021), 41 rows were taken and one row without a structure identifier was excluded, leaving 40 measured features. The 107 GlyTouCan identifiers assigned to these rows were converted to WURCS (107/107) and canonicalized as in Methods 2.2. Two sources of multiple candidates were kept rather than resolved: 24 rows carry several identifiers (up to 13), and 30 of the 107 WURCS strings list alternative linkages, of which 18 leave the parent residue ambiguous and were expanded into all concrete structures before canonicalization (e.g. 729 combinations reduce to 304 topologies, 256 to 13, 8 to 2). This gives 823 feature–candidate pairs (822 distinct topologies). Scores are $\log_2$ ratios of the reported condition means with the same $\epsilon$ as above. One mapping (60 enzymes, 22 distinct glycan sets, set sizes 1–29) serves all seven perturbations; only the ranking differs. Target sets and expected directions were listed before the analysis from the deleted reactions; SLC35C1, a GDP-fucose transporter without an enzyme rule, is read through the FUT8 glycan set. No correction is pooled across perturbations.

#### S6. Implementation and retained outputs

GlyEnzEA runs in R $\geq$ 4.2 with igraph, tidyverse, fgsea, fedup and qvalue. Canonical topologies are cached for reuse; canonical structures, incidence matrices, glycan-set equivalence classes and enrichment results are saved.

**Table S1:** Enzyme-centered glycan sets of the Mgat3-knockout mouse glycome: composition, preranked GSEA, correction, degree-preserving (curveball) null and minSize sensitivity. NES and nominal $P$ from fgsea; Storey q is the primary correction and BH the Benjamini–Hochberg adjustment within the family of 16 sets; curveball values are from 1,000 degree-preserving randomizations; the last two columns are the fgsea $P$ with minSize 2 and 3 (sets below the threshold are excluded, shown as NA).

| Glycan set | Member enzymes | Class | Set size | NES | Nominal P | Storey q | BH P | Correction | Curveball NES | Curveball P | P (minSize 2) | P (minSize 3) |
|---|---|---|---|---|---|---|---|---|---|---|---|---|
| MGAT3 | MGAT3 | GT | 7 | -1.6 | 0.003 | 0.033 | 0.049 | Storey | -1.8 | 0.001 | 0.0027 | 0.0027 |
| ST3GAL/ST6GAL/CMAS | CMAS, ST3GAL1, ST3GAL3, ST3GAL4, ST3GAL6, ST6GAL1, ST6GAL2 | GT/Other | 12 | 1.44 | 0.074 | 0.32 | 0.47 | Storey | 1.44 | 0.026 | 0.081 | 0.081 |
| FUCA1 | FUCA1 | GH | 8 | -1.32 | 0.12 | 0.32 | 0.47 | Storey | -1.46 | 0.029 | 0.12 | 0.12 |
| FUT4/7/9 | FUT4, FUT7, FUT9 | GT | 23 | 1.25 | 0.12 | 0.32 | 0.47 | Storey | 1.16 | 0.047 | 0.13 | 0.13 |
| B3GALT5 | B3GALT5 | GT | 22 | 1.17 | 0.2 | 0.32 | 0.47 | Storey | 1.09 | 0.065 | 0.2 | 0.2 |
| FUT3/5/6 | FUT3, FUT5, FUT6 | GT | 22 | 1.16 | 0.21 | 0.32 | 0.47 | Storey | 1.08 | 0.079 | 0.21 | 0.21 |
| B4GALT/NEU | B3GALT2, B4GALT1, B4GALT2, B4GALT3, B4GALT4, B4GALT5, NEU1, NEU2, NEU3, NEU4 | GH/GT | 24 | 1.15 | 0.23 | 0.32 | 0.47 | Storey | 1.05 | 0.086 | 0.24 | 0.24 |
| MAN1A1/MAN1A2/MAN1C1 | MAN1A1, MAN1A2, MAN1C1 | GH | 7 | -1.2 | 0.24 | 0.32 | 0.47 | Storey | -1.3 | 0.093 | 0.24 | 0.24 |
| MGAT1/2/5B | MGAT1, MGAT2, MGAT5B | GT | 29 | -1.22 | 0.31 | 0.36 | 0.55 | Storey | -1.25 | 0.2 | 0.3 | 0.3 |
| ALG11/12/3/9 | ALG11, ALG12, ALG3, ALG9 | GT | 8 | -1.03 | 0.45 | 0.48 | 0.73 | Storey | -1.13 | 0.2 | 0.45 | 0.45 |
| MANEA | MANEA | Other | 1 | -0.91 | 0.65 | 0.56 | 0.85 | Storey | -0.85 | 0.39 | NA | NA |
| MAN2A1/2 | MAN2A1, MAN2A2 | GH | 26 | -0.82 | 0.72 | 0.56 | 0.85 | Storey | -0.84 | 0.57 | 0.72 | 0.72 |
| MGAT4/5 | MGAT4A, MGAT4B, MGAT4C, MGAT4D, MGAT5 | GT/Other | 16 | -0.82 | 0.74 | 0.56 | 0.85 | Storey | -0.89 | 0.48 | 0.74 | 0.74 |
| FUT8 | FUT8 | GT | 28 | -0.81 | 0.77 | 0.56 | 0.85 | Storey | -0.84 | 0.57 | 0.77 | 0.77 |
| MAN1B1 | MAN1B1 | GH | 2 | -0.81 | 0.79 | 0.56 | 0.85 | Storey | -0.85 | 0.36 | 0.79 | NA |
| FUT1/2 | FUT1, FUT2 | GT | 9 | 0.57 | 0.98 | 0.65 | 0.98 | Storey | 0.61 | 0.29 | 0.98 | 0.98 |

**Table S2:** Preranked GSEA of all 22 enzyme-centered glycan sets in each of the seven HEK293 perturbations (Huang et al., 2021). Target set marks the set expected to change from the deleted reactions. q is the within-perturbation value used in the paper: the Storey q-value, or the BH adjustment (BH*) when $\pi_0$ could not be estimated, as stated in the last column; BH $P$ is the BH-adjusted $P$ for every set.

| Perturbation | Glycan set | Target set | Set size | NES | Nominal P | q | BH P | Correction |
|---|---|---|---|---|---|---|---|---|
| B4GALNT3&4-KO | CHST9/8 | | 2 | -1.42 | 0.0024 | 0.035 | 0.053 | Storey |
| B4GALNT3&4-KO | B3GALNT2/ B4GALNT3/B4GALNT4 | yes | 3 | -1.44 | 0.016 | 0.1 | 0.16 | Storey |
| B4GALNT3&4-KO | MGAT3 | | 5 | 1.45 | 0.022 | 0.1 | 0.16 | Storey |
| B4GALNT3&4-KO | CMAS | | 18 | 1.23 | 0.19 | 0.44 | 0.66 | Storey |
| B4GALNT3&4-KO | ST3GAL3/ST3GAL4/ ST3GAL6/ST6GAL1/ ST6GAL2/ST3GAL1 | | 17 | 1.23 | 0.2 | 0.44 | 0.66 | Storey |
| B4GALNT3&4-KO | MGAT1/2/5B | | 29 | 1.07 | 0.31 | 0.44 | 0.66 | Storey |
| B4GALNT3&4-KO | B3GALT5 | | 20 | -1.09 | 0.33 | 0.44 | 0.66 | Storey |
| B4GALNT3&4-KO | MGAT4/5 | | 16 | -1.1 | 0.34 | 0.44 | 0.66 | Storey |
| B4GALNT3&4-KO | MAN2A1/2 | | 28 | 1.04 | 0.37 | 0.44 | 0.66 | Storey |
| B4GALNT3&4-KO | FUT1/2 | | 2 | -1.14 | 0.38 | 0.44 | 0.66 | Storey |
| B4GALNT3&4-KO | FUT8 | | 21 | -1.05 | 0.38 | 0.44 | 0.66 | Storey |
| B4GALNT3&4-KO | FUT3/4/9 | | 15 | -1.07 | 0.41 | 0.44 | 0.66 | Storey |
| B4GALNT3&4-KO | FUT5/6 | | 14 | -1.07 | 0.42 | 0.44 | 0.66 | Storey |
| B4GALNT3&4-KO | FUT7 | | 14 | -1.07 | 0.42 | 0.44 | 0.66 | Storey |
| B4GALNT3&4-KO | B4GALT/NEU | | 24 | -0.97 | 0.54 | 0.53 | 0.79 | Storey |
| B4GALNT3&4-KO | MAN1B1 | | 1 | -0.92 | 0.6 | 0.55 | 0.83 | Storey |
| B4GALNT3&4-KO | MANEA | | 2 | -0.9 | 0.65 | 0.56 | 0.84 | Storey |
| B4GALNT3&4-KO | FUCA1 | | 20 | 0.84 | 0.71 | 0.56 | 0.85 | Storey |
| B4GALNT3&4-KO | ALG11/12/3/9 | | 9 | 0.77 | 0.74 | 0.56 | 0.85 | Storey |
| B4GALNT3&4-KO | ALG5/ALG6/UGGT1/ UGGT2 | | 1 | -0.81 | 0.8 | 0.58 | 0.88 | Storey |
| B4GALNT3&4-KO | MAN1A1/MAN1A2/ MAN1C1 | | 5 | -0.48 | 0.94 | 0.65 | 0.99 | Storey |
| B4GALNT3&4-KO | B3GNT2/4/8 | | 1 | -0.68 | 1 | 0.66 | 1 | Storey |
| MGAT1-KO | FUT8 | | 21 | -1.86 | $7 \times 10^{-4}$ | 0.015 | 0.015 | BH* |
| MGAT1-KO | MGAT1/2/5B | yes | 29 | -1.82 | 0.0026 | 0.029 | 0.029 | BH* |
| MGAT1-KO | MAN2A1/2 | | 28 | -1.68 | 0.0098 | 0.072 | 0.072 | BH* |
| MGAT1-KO | B4GALT/NEU | | 24 | -1.56 | 0.021 | 0.11 | 0.11 | BH* |
| MGAT1-KO | B3GALT5 | | 20 | -1.4 | 0.06 | 0.26 | 0.26 | BH* |
| MGAT1-KO | MGAT4/5 | | 16 | -1.37 | 0.073 | 0.27 | 0.27 | BH* |
| MGAT1-KO | FUT5/6 | | 14 | -1.31 | 0.11 | 0.35 | 0.35 | BH* |
| MGAT1-KO | FUT3/4/9 | | 15 | -1.26 | 0.14 | 0.36 | 0.36 | BH* |
| MGAT1-KO | CMAS | | 18 | -1.24 | 0.17 | 0.36 | 0.36 | BH* |
| MGAT1-KO | MGAT3 | | 5 | -1.23 | 0.18 | 0.36 | 0.36 | BH* |
| MGAT1-KO | FUT7 | | 14 | -1.2 | 0.21 | 0.36 | 0.36 | BH* |
| MGAT1-KO | MANEA | | 2 | 1.19 | 0.22 | 0.36 | 0.36 | BH* |
| MGAT1-KO | FUCA1 | | 20 | 1.15 | 0.24 | 0.36 | 0.36 | BH* |
| MGAT1-KO | ST3GAL3/ST3GAL4/ ST3GAL6/ST6GAL1/ ST6GAL2/ST3GAL1 | | 17 | -1.17 | 0.24 | 0.36 | 0.36 | BH* |
| MGAT1-KO | ALG11/12/3/9 | | 9 | 1.18 | 0.25 | 0.36 | 0.36 | BH* |
| MGAT1-KO | FUT1/2 | | 2 | -1.09 | 0.35 | 0.45 | 0.45 | BH* |
| MGAT1-KO | MAN1B1 | | 1 | 1.12 | 0.35 | 0.45 | 0.45 | BH* |
| MGAT1-KO | B3GNT2/4/8 | | 1 | -1.08 | 0.4 | 0.49 | 0.49 | BH* |
| MGAT1-KO | ALG5/ALG6/UGGT1/ UGGT2 | | 1 | 0.95 | 0.6 | 0.69 | 0.69 | BH* |
| MGAT1-KO | MAN1A1/MAN1A2/ MAN1C1 | | 5 | 0.89 | 0.66 | 0.72 | 0.72 | BH* |
| MGAT1-KO | B3GALNT2/ B4GALNT3/B4GALNT4 | | 3 | -0.73 | 0.89 | 0.9 | 0.9 | BH* |
| MGAT1-KO | CHST9/8 | | 2 | -0.73 | 0.9 | 0.9 | 0.9 | BH* |
| MGAT2-KO | MGAT4/5 | | 16 | -1.8 | $1.5 \times 10^{-5}$ | 0.00021 | 0.00033 | Storey |
| MGAT2-KO | B3GALT5 | | 20 | -1.69 | 0.0011 | 0.0078 | 0.012 | Storey |
| MGAT2-KO | FUT8 | | 21 | -1.55 | 0.011 | 0.052 | 0.081 | Storey |
| MGAT2-KO | FUT5/6 | | 14 | -1.47 | 0.02 | 0.07 | 0.11 | Storey |
| MGAT2-KO | MAN2A1/2 | | 28 | -1.48 | 0.031 | 0.081 | 0.13 | Storey |
| MGAT2-KO | MGAT1/2/5B | yes | 29 | -1.47 | 0.034 | 0.081 | 0.13 | Storey |
| MGAT2-KO | B4GALT/NEU | | 24 | -1.4 | 0.055 | 0.11 | 0.17 | Storey |
| MGAT2-KO | FUT3/4/9 | | 15 | -1.37 | 0.061 | 0.11 | 0.17 | Storey |
| MGAT2-KO | MGAT3 | | 5 | -1.22 | 0.15 | 0.19 | 0.29 | Storey |
| MGAT2-KO | B3GNT2/4/8 | | 1 | -1.25 | 0.15 | 0.19 | 0.29 | Storey |
| MGAT2-KO | FUT7 | | 14 | -1.25 | 0.15 | 0.19 | 0.29 | Storey |
| MGAT2-KO | FUT1/2 | | 2 | -1.22 | 0.16 | 0.19 | 0.29 | Storey |
| MGAT2-KO | CMAS | | 18 | -1.11 | 0.32 | 0.35 | 0.55 | Storey |
| MGAT2-KO | ALG11/12/3/9 | | 9 | 1.04 | 0.35 | 0.36 | 0.56 | Storey |
| MGAT2-KO | CHST9/8 | | 2 | -1.04 | 0.48 | 0.45 | 0.7 | Storey |
| MGAT2-KO | ST3GAL3/ST3GAL4/ ST3GAL6/ST6GAL1/ ST6GAL2/ST3GAL1 | | 17 | -0.97 | 0.53 | 0.47 | 0.73 | Storey |
| MGAT2-KO | MAN1A1/MAN1A2/ MAN1C1 | | 5 | 0.75 | 0.75 | 0.61 | 0.96 | Storey |
| MGAT2-KO | ALG5/ALG6/UGGT1/ UGGT2 | | 1 | -0.78 | 0.85 | 0.61 | 0.96 | Storey |
| MGAT2-KO | FUCA1 | | 20 | 0.81 | 0.87 | 0.61 | 0.96 | Storey |
| MGAT2-KO | B3GALNT2/ B4GALNT3/B4GALNT4 | | 3 | -0.74 | 0.87 | 0.61 | 0.96 | Storey |

*continued on next page*

**Table S2:** (continued)

| Perturbation | Glycan set | Target set | Set size | NES | Nominal P | q | BH P | Correction |
|---|---|---|---|---|---|---|---|---|
| MGAT2-KO | MAN1B1 | | 1 | -0.71 | 0.95 | 0.64 | 0.99 | Storey |
| MGAT2-KO | MANEA | | 2 | 0 | 1 | 0.64 | 1 | Storey |
| MGAT4A&4B-KO | MGAT1/2/5B | | 29 | 1.28 | 0.096 | 0.9 | 0.9 | BH* |
| MGAT4A&4B-KO | CHST9/8 | | 2 | -1.28 | 0.21 | 0.9 | 0.9 | BH* |
| MGAT4A&4B-KO | MGAT3 | | 5 | 1.28 | 0.24 | 0.9 | 0.9 | BH* |
| MGAT4A&4B-KO | MAN2A1/2 | | 28 | 1.11 | 0.29 | 0.9 | 0.9 | BH* |
| MGAT4A&4B-KO | B3GNT2/4/8 | | 1 | 1.09 | 0.4 | 0.9 | 0.9 | BH* |
| MGAT4A&4B-KO | B3GALNT2/ B4GALNT3/B4GALNT4 | | 3 | -1.15 | 0.42 | 0.9 | 0.9 | BH* |
| MGAT4A&4B-KO | ST3GAL3/ST3GAL4/ ST3GAL6/ST6GAL1/ ST6GAL2/ST3GAL1 | | 17 | 1.06 | 0.44 | 0.9 | 0.9 | BH* |
| MGAT4A&4B-KO | CMAS | | 18 | 1.05 | 0.45 | 0.9 | 0.9 | BH* |
| MGAT4A&4B-KO | MANEA | | 2 | -1 | 0.55 | 0.9 | 0.9 | BH* |
| MGAT4A&4B-KO | ALG5/ALG6/UGGT1/ UGGT2 | | 1 | -0.92 | 0.6 | 0.9 | 0.9 | BH* |
| MGAT4A&4B-KO | ALG11/12/3/9 | | 9 | 0.83 | 0.64 | 0.9 | 0.9 | BH* |
| MGAT4A&4B-KO | B4GALT/NEU | | 24 | 0.87 | 0.67 | 0.9 | 0.9 | BH* |
| MGAT4A&4B-KO | B3GALT5 | | 20 | -0.9 | 0.72 | 0.9 | 0.9 | BH* |
| MGAT4A&4B-KO | FUCA1 | | 20 | 0.76 | 0.78 | 0.9 | 0.9 | BH* |
| MGAT4A&4B-KO | MGAT4/5 | yes | 16 | -0.87 | 0.8 | 0.9 | 0.9 | BH* |
| MGAT4A&4B-KO | MAN1B1 | | 1 | -0.81 | 0.8 | 0.9 | 0.9 | BH* |
| MGAT4A&4B-KO | FUT8 | | 21 | -0.86 | 0.82 | 0.9 | 0.9 | BH* |
| MGAT4A&4B-KO | FUT5/6 | | 14 | -0.86 | 0.83 | 0.9 | 0.9 | BH* |
| MGAT4A&4B-KO | MAN1A1/MAN1A2/ MAN1C1 | | 5 | 0.68 | 0.86 | 0.9 | 0.9 | BH* |
| MGAT4A&4B-KO | FUT3/4/9 | | 15 | -0.83 | 0.86 | 0.9 | 0.9 | BH* |
| MGAT4A&4B-KO | FUT7 | | 14 | -0.83 | 0.86 | 0.9 | 0.9 | BH* |
| MGAT4A&4B-KO | FUT1/2 | | 2 | -0.6 | 0.92 | 0.92 | 0.92 | BH* |
| MGAT5-KO | MAN2A1/2 | | 28 | 1.29 | 0.11 | 0.64 | 0.64 | BH* |
| MGAT5-KO | MGAT1/2/5B | | 29 | 1.25 | 0.13 | 0.64 | 0.64 | BH* |
| MGAT5-KO | CHST9/8 | | 2 | 1.3 | 0.15 | 0.64 | 0.64 | BH* |
| MGAT5-KO | B3GNT2/4/8 | | 1 | -1.25 | 0.15 | 0.64 | 0.64 | BH* |
| MGAT5-KO | FUT1/2 | | 2 | -1.36 | 0.15 | 0.64 | 0.64 | BH* |
| MGAT5-KO | MANEA | | 2 | -1.31 | 0.2 | 0.64 | 0.64 | BH* |
| MGAT5-KO | ALG5/ALG6/UGGT1/ UGGT2 | | 1 | -1.19 | 0.25 | 0.64 | 0.64 | BH* |
| MGAT5-KO | FUT8 | | 21 | 1.13 | 0.29 | 0.64 | 0.64 | BH* |
| MGAT5-KO | MGAT3 | | 5 | 1.16 | 0.36 | 0.64 | 0.64 | BH* |
| MGAT5-KO | B3GALT5 | | 20 | -1.04 | 0.4 | 0.64 | 0.64 | BH* |
| MGAT5-KO | B3GALNT2/ B4GALNT3/B4GALNT4 | | 3 | 1.08 | 0.4 | 0.64 | 0.64 | BH* |
| MGAT5-KO | MAN1A1/MAN1A2/ MAN1C1 | | 5 | -1.13 | 0.42 | 0.64 | 0.64 | BH* |
| MGAT5-KO | ST3GAL3/ST3GAL4/ ST3GAL6/ST6GAL1/ ST6GAL2/ST3GAL1 | | 17 | -1.03 | 0.43 | 0.64 | 0.64 | BH* |
| MGAT5-KO | ALG11/12/3/9 | | 9 | -1.08 | 0.44 | 0.64 | 0.64 | BH* |
| MGAT5-KO | CMAS | | 18 | -1.02 | 0.44 | 0.64 | 0.64 | BH* |
| MGAT5-KO | MGAT4/5 | yes | 16 | -0.98 | 0.52 | 0.71 | 0.71 | BH* |
| MGAT5-KO | MAN1B1 | | 1 | -0.92 | 0.65 | 0.79 | 0.79 | BH* |
| MGAT5-KO | B4GALT/NEU | | 24 | 0.88 | 0.69 | 0.79 | 0.79 | BH* |
| MGAT5-KO | FUT7 | | 14 | 0.8 | 0.74 | 0.79 | 0.79 | BH* |
| MGAT5-KO | FUT5/6 | | 14 | 0.79 | 0.75 | 0.79 | 0.79 | BH* |
| MGAT5-KO | FUT3/4/9 | | 15 | 0.79 | 0.75 | 0.79 | 0.79 | BH* |
| MGAT5-KO | FUCA1 | | 20 | 0.66 | 0.94 | 0.94 | 0.94 | BH* |
| SLC35C1-KO | B4GALT/NEU | | 24 | -1.68 | 0.003 | 0.066 | 0.066 | BH* |
| SLC35C1-KO | B3GALT5 | | 20 | -1.57 | 0.009 | 0.071 | 0.071 | BH* |
| SLC35C1-KO | FUT8 | yes | 21 | -1.56 | 0.011 | 0.071 | 0.071 | BH* |
| SLC35C1-KO | MGAT1/2/5B | | 29 | -1.61 | 0.013 | 0.071 | 0.071 | BH* |
| SLC35C1-KO | FUT5/6 | | 14 | -1.4 | 0.049 | 0.18 | 0.18 | BH* |
| SLC35C1-KO | FUT3/4/9 | | 15 | -1.38 | 0.059 | 0.18 | 0.18 | BH* |
| SLC35C1-KO | MAN2A1/2 | | 28 | -1.42 | 0.061 | 0.18 | 0.18 | BH* |
| SLC35C1-KO | CMAS | | 18 | -1.37 | 0.066 | 0.18 | 0.18 | BH* |
| SLC35C1-KO | MGAT4/5 | | 16 | -1.33 | 0.088 | 0.22 | 0.22 | BH* |
| SLC35C1-KO | FUT7 | | 14 | -1.29 | 0.12 | 0.26 | 0.26 | BH* |
| SLC35C1-KO | ST3GAL3/ST3GAL4/ ST3GAL6/ST6GAL1/ ST6GAL2/ST3GAL1 | | 17 | -1.26 | 0.15 | 0.29 | 0.29 | BH* |
| SLC35C1-KO | ALG11/12/3/9 | | 9 | -1.21 | 0.19 | 0.32 | 0.32 | BH* |
| SLC35C1-KO | FUCA1 | | 20 | 1.12 | 0.19 | 0.32 | 0.32 | BH* |
| SLC35C1-KO | FUT1/2 | | 2 | -1.19 | 0.2 | 0.32 | 0.32 | BH* |
| SLC35C1-KO | MAN1A1/MAN1A2/ MAN1C1 | | 5 | -1.13 | 0.27 | 0.4 | 0.4 | BH* |
| SLC35C1-KO | B3GALNT2/ B4GALNT3/B4GALNT4 | | 3 | -1.06 | 0.39 | 0.54 | 0.54 | BH* |
| SLC35C1-KO | CHST9/8 | | 2 | -1.04 | 0.44 | 0.55 | 0.55 | BH* |
| SLC35C1-KO | MANEA | | 2 | 1.06 | 0.45 | 0.55 | 0.55 | BH* |
| SLC35C1-KO | ALG5/ALG6/UGGT1/ UGGT2 | | 1 | 1.02 | 0.5 | 0.58 | 0.58 | BH* |
| SLC35C1-KO | MGAT3 | | 5 | -0.86 | 0.73 | 0.8 | 0.8 | BH* |
| SLC35C1-KO | MAN1B1 | | 1 | 0.81 | 0.8 | 0.8 | 0.8 | BH* |
| SLC35C1-KO | B3GNT2/4/8 | | 1 | -0.81 | 0.8 | 0.8 | 0.8 | BH* |
| T-KO (MAN1A1&A2&B1) | FUT8 | | 21 | -1.94 | $6.2 \times 10^{-5}$ | 0.0014 | 0.0014 | BH* |

*continued on next page*

**Table S2:** (continued)

| Perturbation | Glycan set | Target set | Set size | NES | Nominal P | q | BH P | Correction |
|---|---|---|---|---|---|---|---|---|
| T-KO (MAN1A1&A2&B1) | B3GALT5 | | 20 | -1.78 | 0.0014 | 0.011 | 0.011 | BH* |
| T-KO (MAN1A1&A2&B1) | ALG11/12/3/9 | | 9 | 2.27 | 0.0015 | 0.011 | 0.011 | BH* |
| T-KO (MAN1A1&A2&B1) | MAN2A1/2 | | 28 | -1.71 | 0.006 | 0.033 | 0.033 | BH* |
| T-KO (MAN1A1&A2&B1) | FUT5/6 | | 14 | -1.58 | 0.011 | 0.038 | 0.038 | BH* |
| T-KO (MAN1A1&A2&B1) | MGAT1/2/5B | | 29 | -1.65 | 0.012 | 0.038 | 0.038 | BH* |
| T-KO (MAN1A1&A2&B1) | FUCA1 | | 20 | 1.71 | 0.012 | 0.038 | 0.038 | BH* |
| T-KO (MAN1A1&A2&B1) | CMAS | | 18 | -1.56 | 0.016 | 0.041 | 0.041 | BH* |
| T-KO (MAN1A1&A2&B1) | MGAT4/5 | | 16 | -1.54 | 0.018 | 0.041 | 0.041 | BH* |
| T-KO (MAN1A1&A2&B1) | MAN1A1/MAN1A2/ MAN1C1 | yes | 5 | 1.7 | 0.02 | 0.041 | 0.041 | BH* |
| T-KO (MAN1A1&A2&B1) | FUT3/4/9 | | 15 | -1.53 | 0.02 | 0.041 | 0.041 | BH* |
| T-KO (MAN1A1&A2&B1) | B4GALT/NEU | | 24 | -1.51 | 0.031 | 0.056 | 0.056 | BH* |
| T-KO (MAN1A1&A2&B1) | FUT7 | | 14 | -1.46 | 0.039 | 0.066 | 0.066 | BH* |
| T-KO (MAN1A1&A2&B1) | ST3GAL3/ST3GAL4/ ST3GAL6/ST6GAL1/ ST6GAL2/ST3GAL1 | | 17 | -1.42 | 0.056 | 0.087 | 0.087 | BH* |
| T-KO (MAN1A1&A2&B1) | MGAT3 | | 5 | -1.31 | 0.11 | 0.16 | 0.16 | BH* |
| T-KO (MAN1A1&A2&B1) | B3GNT2/4/8 | | 1 | -1.22 | 0.2 | 0.27 | 0.27 | BH* |
| T-KO (MAN1A1&A2&B1) | MAN1B1 | | 1 | 1.19 | 0.25 | 0.32 | 0.32 | BH* |
| T-KO (MAN1A1&A2&B1) | CHST9/8 | | 2 | -1.04 | 0.43 | 0.52 | 0.52 | BH* |
| T-KO (MAN1A1&A2&B1) | MANEA | | 2 | 1.02 | 0.45 | 0.52 | 0.52 | BH* |
| T-KO (MAN1A1&A2&B1) | B3GALNT2/ B4GALNT3/B4GALNT4 | | 3 | -0.99 | 0.51 | 0.56 | 0.56 | BH* |
| T-KO (MAN1A1&A2&B1) | ALG5/ALG6/UGGT1/ UGGT2 | | 1 | 0.81 | 0.8 | 0.84 | 0.84 | BH* |
| T-KO (MAN1A1&A2&B1) | FUT1/2 | | 2 | -0.76 | 0.87 | 0.87 | 0.87 | BH* |

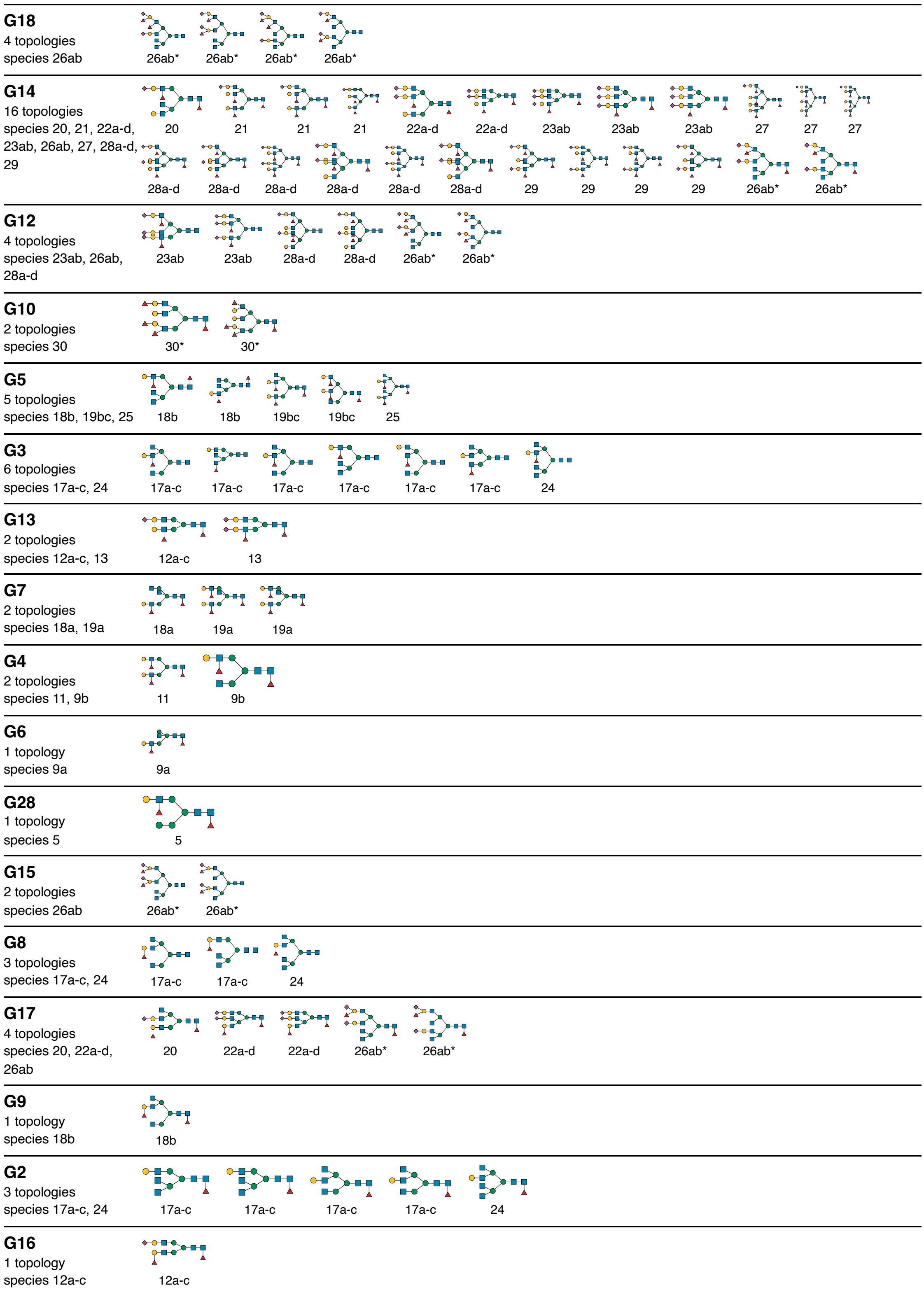


**Figure S1:** Candidate structures in each display column of Figure 3. One row per column of Figure 3 (G labels), in the order of Figure 3 from left to right; each row shows the candidate structures whose topology has that enzyme-mapping pattern, drawn in SNFG without linkage labels, with the measured features (numbering of Nakano et al., 2019) to which each candidate belongs. Several candidate structures can share one topology because linkage positions are not retained, so a row can show more drawings than topologies. Candidates of features 6, 26ab and 30 (asterisks) were entered manually without GlyTouCan identifiers and are drawn from their topology. (page 1 of 2)

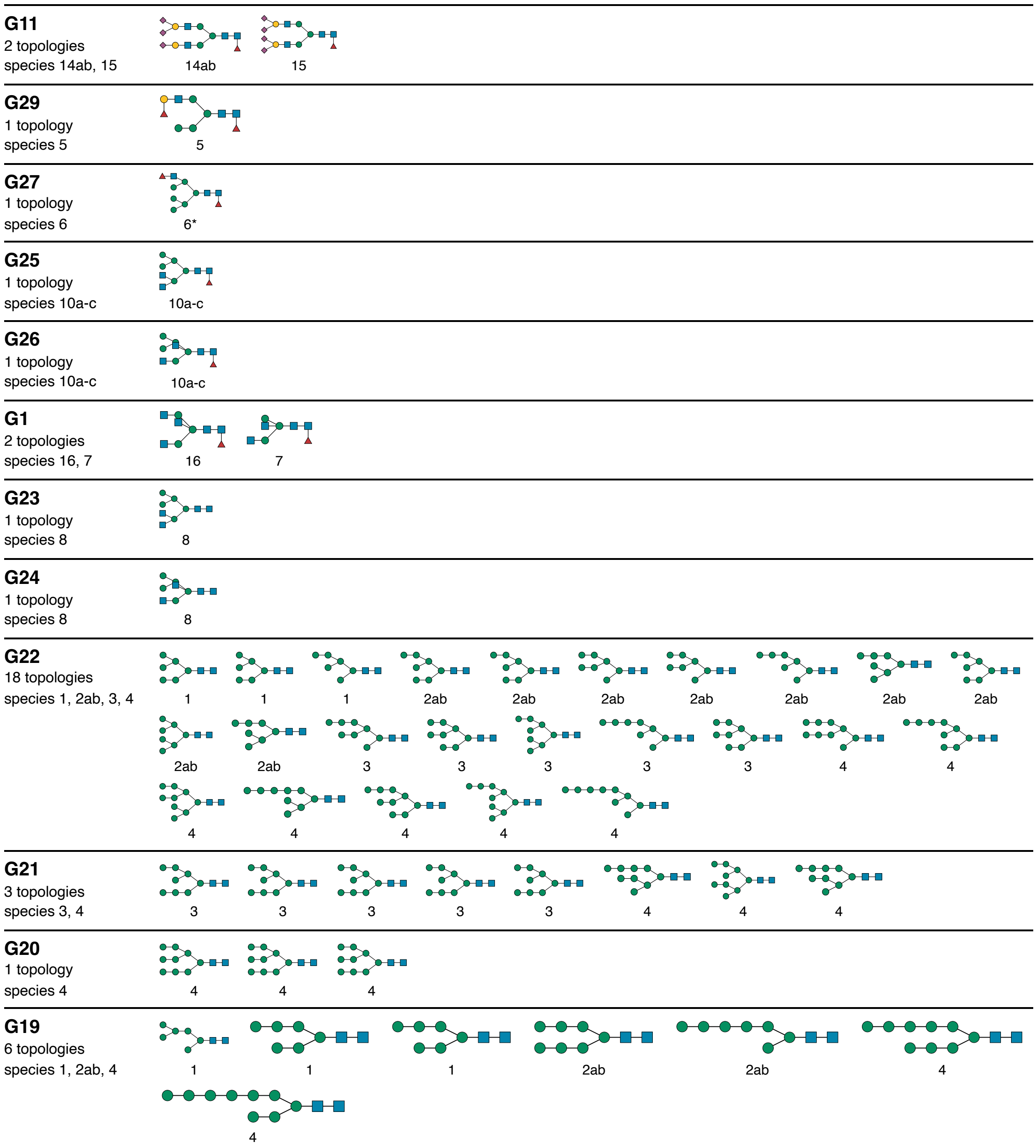


**Figure S1** (continued, page 2 of 2).

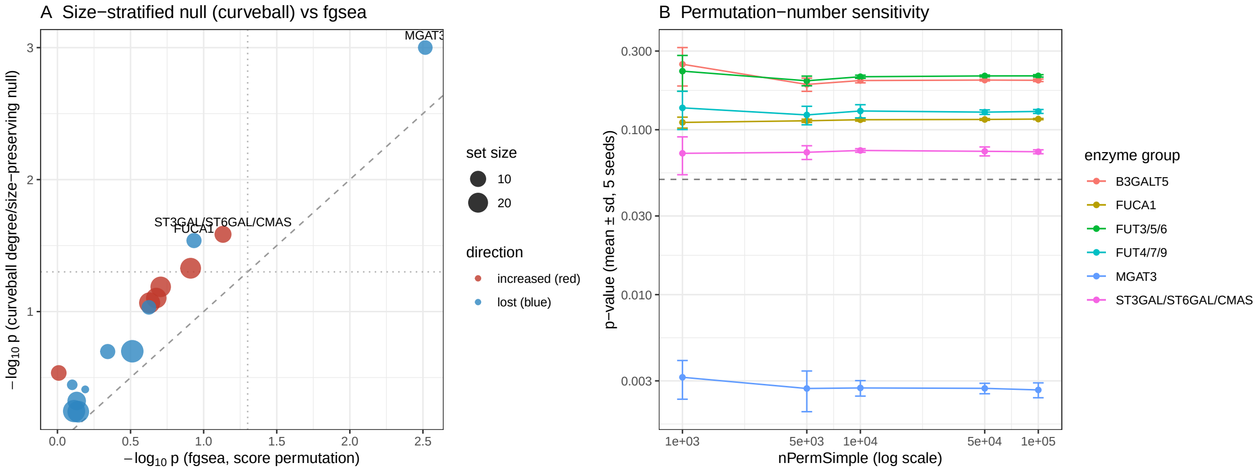


**Figure S2:** Null checks for the mouse enrichment results. (A) fgsea score-permutation $P$ against the empirical $P$ from 1,000 degree-preserving curveball randomizations of the enzyme–feature matrix, both $-\log_{10}$; point size is set size, colour the direction of NES; dashed line $y = x$, dotted lines $P = 0.05$. The MGAT3 glycan set is the most extreme set under both nulls and reaches the empirical resolution of the curveball null ($P = 0.001$). (B) fgsea $P$ (mean $\pm$ SD over five seeds) for nPermSimple from $10^3$ to $10^5$; the dashed line marks $P = 0.05$. Point estimates are stable; for the MGAT3 glycan set the Storey q-value is below 0.05 in every run.

#### S7. Use of generative AI and author contributions

Generative AI was used for English-language editing, translation and assistance with code writing. Codex (OpenAI) also assisted manuscript formatting and submission preparation. AI assistance is disclosed here as part of the research and writing workflow; responsibility for the manuscript and code rests with the authors.

##### Author contributions (CRediT)

Koichi Himori: methodology, software, formal analysis, validation and writing—original draft.

Bingyuan Zhang: methodology and formal analysis.

Yusuke Matsui: conceptualization, methodology, validation, writing—original draft and supervision.
